# Simultaneous quantification of *Vibrio metoecus* and *Vibrio cholerae* with its O1 serogroup and toxigenic subpopulations in environmental reservoirs

**DOI:** 10.1101/822551

**Authors:** Tania Nasreen, Nora A. S. Hussain, Mohammad Tarequl Islam, Fabini D. Orata, Paul C. Kirchberger, Munirul Alam, Stephanie K. Yanow, Yann F. Boucher

**Affiliations:** Department of Biological Sciences, University of Alberta, Edmonton, Alberta T6G 2E9, Canada; Department of Chemical and Materials Engineering, University of Alberta, Edmonton, Alberta T6G 1H9, Canada; Department of Integrative Biology, University of Texas at Austin, Austin, Texas 78712, USA; Centre for Communicable Diseases, International Centre for Diarrhoeal Disease Research, Bangladesh (ICDDR, B), Dhaka, Bangladesh; School of Public Health, University of Alberta, Edmonton, Alberta T6G 2E9, Canada; Department of Medical Microbiology and Immunology, University of Alberta, Edmonton, Alberta T6G 2E9, Canada

**Author notes:** Corresponding Author: Yann F. Boucher.

**Keywords:** *Vibrio cholerae*, *Vibrio metoecus*, qPCR, serogroup, toxigenic, non-toxigenic

## Abstract

*Vibrio metoecus* is a recently described and little studied causative agent of opportunistic infections in humans, often coexisting with *V. cholerae* in aquatic environments. However, the relative abundance of *V. metoecus* with *V. cholera*e and their population dynamics in aquatic reservoirs is still unknown. We developed a multiplex qPCR assay with a limit of detection of three copies per reaction to simultaneously quantify total *V. metoecus* and *V. cholerae* abundance, as well as the toxigenic and O1 serogroup subpopulations of *V. cholerae* from environmental samples. Four different genes were targeted as specific markers for individual *Vibrio* species or subpopulations; *viuB*, a gene encoding a vibriobactin utilization protein, was used to quantify the total *V. cholerae* population. The cholera toxin gene *ctxA* provided an estimation of toxigenic *V. cholerae* abundance, while the *rfbO1* gene specifically detected and quantified *V. cholerae* belonging to the O1 serogroup, which includes almost all lineages of the species responsible for the majority of past and ongoing cholera pandemics. To measure *V. metoecus* abundance, the gene *mcp*, encoding methyl accepting chemotaxis protein, was used. Marker specificity was confirmed by testing several isolates of *V. cholerae* and *V. metoecus* alongside negative controls of isolates within and outside of the *Vibrio* genus. Analysis of environmental water samples collected from four different geographic locations including cholera-endemic (Dhaka, Kuakata and Mathbaria in Bangladesh) and non-endemic (Oyster Pond in Falmouth, Massachusetts, USA) regions showed that *V. metoecus* was only present in the USA site, recurring seasonally. Within the coastal USA site, the non-toxigenic O1 serogroup represented up to ∼18% of the total *V. cholerae* population. *V. cholerae* toxigenic O1 serogroup was absent or present in low abundance in coastal Bangladesh (Kuakata and Mathbaria) but constituted a relatively high proportion of the total *V. cholerae* population sustained throughout the year in inland Bangladesh (Dhaka). A preference for host/particle attachment was observed, as the majority of cells from both *Vibrio* species (>90%) were identified in the largest water size fraction sampled, composed of particles or organisms >63 μm and their attached bacteria. This is the first study to apply a culture-independent method to quantify *V. cholerae* or *V. metoecus* directly in environmental reservoirs of areas endemic and non-endemic for cholera on significant temporal and spatial scales.

**SIGNIFICANCE:** Cholera is a life-threatening disease that requires immediate intervention; it is of prime importance to have fast, accurate and sensitive means to detect *V. cholerae*. Consistent environmental monitoring of the abundance of *V. cholerae* along with its toxigenic and O1 serogroup subpopulations could facilitate the determination of the actual distribution of this organism in aquatic reservoirs and thus help to predict an outbreak before it strikes. The lack of substantial temporal and spatial environmental sampling, along with specific quantitative measures, has made this goal elusive so far. The same is true for *V. metoecus*, a close relative of *V. cholerae* which has been associated with several clinical infections and could likely pose an emerging threat, readily exchanging genetic material with its more famous relative.

## INTRODUCTION

*Vibrio cholerae* is an autochthonous aquatic bacterium (Colwell et al., 1977) which shows variable physiologies, from non-pathogenic to extremely virulent strains capable of causing a life-threatening diarrheal infection, cholera (Alam et al., 2006). According to the World Health Organization (2016), every year, 1.4 to 4 million people are infected with cholera, and 21,000 to 143,000 people die from this disease (WHO, 2016). This species comprises over 200 serogroups (Thompson et al., 2004), but strains of the O1 and O139 serogroups are distinguished as the most virulent, has caused some of the most devastating pandemics in human history (Ali et al., 2012; Chun et al., 2009; Islam et al., 1994; Kaper et al., 1995). Other serogroups are collectively known as non-O1/non-O139 and can cause sporadic outbreaks of diarrheal disease not severe like cholera (Dutta et al., 2013). *V. cholerae* O139 was only found to be associated with isolated cholera cases after two epidemics occurred in 1993 and in 2002 (Faruque et al., 2003; Ghosh et al., 2016). During 2005 in Bangladesh O139 serogroup has been isolated sporadically from both clinical and environmental samples but there was not any reported largescale outbreak of cholera caused by *V. cholerae* O139. Further study confirmed that four *V. cholerae* isolated from cholera patients identified as *V. cholerae* O139 during 2013 and 2014 in Bangladesh by typing and whole genome sequencing (Alam et al., 2006; Chowdhury et al., 2015; Rashed et al., 2013). However, *V. cholerae* O1 has been prevalent in the Ganges Delta for several hundred years; this remains the only place in the world where cholera has been continually endemic since the first modern pandemic in 1817 (Boucher et al., 2015; Mutreja et al., 2011). Much research has been done to characterize the physiology of pathogenic *V. cholerae* strains, but still, we have limited understanding of the variation of *V. cholerae* population composition and abundance over time and between areas, which influences where and when epidemics are likely to occur.

*Vibrio metoecus* is a recently described species, which has been co-isolated with *V. cholerae* from aquatic environments in the USA East coast (Choopun, 2004; Haley et al., 2010; Kirchberger et al., 2014). Based on biochemical, genotypic and phylogenetic evidence, *V. metoecus* is the closest known relative of *V. cholerae* (Kirchberger et al., 2014). It has been recovered not only from the environment but also from a variety of human specimens (blood, stool, ear and leg wounds) in opportunistic infections across the USA (CDC, Atlanta, Georgia, USA)(Kirchberger et al., 2014). Currently, no studies have assessed the abundance of *V. metoecus* in aquatic habitats. Information on the distribution of *V. metoecus* in its natural reservoirs is essential not only to provide insight into transmission routes from the environment to human, but also because of its co-occurrence with *V. cholerae* and the occurrence of possible horizontal gene transfer (HGT) between the species, which could be responsible for producing more virulent strains of *V. metoecus* (Orata et al., 2015).

Most surveys of *V. cholerae* used culture-based methods, requiring significant time and effort, and their reliance on selective enrichment prevents any form of absolute quantification (Baron et al., 2007). Moreover, these culture-based techniques are unable to detect the viable but non-culturable (VBNC) form of *V. cholerae* (Colwell et al., 1996; Colwell and Huq, 1994; Miller et al., 2009) and thus underestimate the abundance of toxigenic and non-toxigenic *V. cholerae* in the environment (Huq et al., 1996). To circumvent this problem, several studies have used antibody or hybridization-based molecular detection of toxigenic *V. cholerae* from clinical (Lobitz et al., 2000; Qadri et al., 1995) and environmental samples (Baron et al., 2007; Blackstone et al., 2007; Colwell et al., 1985; Huq et al., 1990; Theron et al., 2000), but most are qualitative, and none have been field-tested to comprehensively survey its abundance in natural environments (Gubala, 2006; Neogi et al., 2010). Another molecular tool, real-time quantitative PCR (qPCR), has been used to measure both the presence and abundance of target species (Abdullah et al., 2018; Haugland et al., 2005). This technique can estimate the presence of as few as three bacterial genome equivalents in a sample and provides the high throughput required to investigate the ecological distribution of the organism (Kralik and Ricchi, 2017). Several gene markers have been developed for the detection of the *V. cholerae* species by qPCR, including the *ompW* gene encoding the outer membrane protein (Bliem et al., 2015), the *hlyA* gene encoding for hemolysin (Bliem et al., 2015; Lyon, 2001) and *gbpA* gene encoding the N-acetyl glucosamine-binding protein A (Vezzulli et al., 2015), and the *rtx* gene cluster encoding the repeats in toxin protein (Lin et al., 1999). During the assessment of different primers used in previous studies, it was found that the commonly used primers designed for *ompW* also amplified that gene from *V. metoecus* (in this study) and *Vibrio mimicus* (Gubala and Proll, 2006). Copy number is also a confounding factor, as *rtx* (Lin et al., 1999), *hlyA* and *ctxA* can be present in multiple copies in *V. cholerae* genomes (Ghosh et al., 2011; Heidelberg et al., 2000). Although *gbpA* is present as a single copy, it can also be found in environmental strains belonging to other *Vibrio* species including *Vibrio alginolyticus, Vibrio metschnikovii, Vibrio mimicus, Vibrio vulnificus* and *Vibrio parahaemolyticus* (Stauder et al., 2012). Lack of species specificity and the presence of multiple copies in single cells make quantitative estimates unreliable. Furthermore, given that a single lineage of *V. cholerae* is responsible for all major outbreaks [phylocore genome/pandemic-generating (PG) *V. cholerae*] (Boucher, 2016; Chun et al., 2009)(FIG S1), detection of other genotypes which are mostly harmless can be misleading in evaluating the risk of outbreaks. Attempts to accurately quantify toxigenic *V. cholerae* using the cholera toxin (CT) gene *ctxA* have been made; while the reference El Tor strain N16961 carries a single copy of this gene, other *V. cholerae* strains carry several copies of this element and it is also occasionally found in other *Vibrio* species (Dalsgaard et al., 2001; Faruque et al., 1997; Heidelberg et al., 2000; Mekalanos et al., 1983). The *rfbO1* gene, targeting *V. cholerae* strains carrying the O1 antigens has also been used to detect pandemic *V. cholerae* (Hoshino et al., 1998; Yamasaki et al., 1996). Although it is a single copy gene, many strains unrelated to this lineage can also carry this gene (Pang et al., 2007).

To overcome the limitations of currently used molecular markers, we developed a multiplex qPCR assay for simultaneous detection and quantification of *V. cholerae* O1 (*rfbO1*), toxigenic *V. cholerae* (*ctxA*), total *V. cholerae* (*viuB*) and *V. metoecus* (*mcp).* This optimized qPCR technique allows the quantitative study of *V. cholerae* populations directly from DNA extracted from aquatic biomass, without the need for cultivation. This overcomes the problem that many *V. cholerae* cells in water are viable but not culturable (VBNC) (Colwell et al., 1985), which has so far made it difficult to the study this organism in the environment difficult. Quantifying the presence of *ctxA* and *rfbO1* simultaneously resolves drawbacks specific to the separate use of these two gene markers. Overestimation of the number of toxigenic cells because of the presence of multiple copies of *ctxA* in single cells is detected as discrepancies with *rfbO1* abundance. Likewise, overestimation of toxigenic O1 cells due to the presence of non-toxigenic O1 strains is identified through discrepancies with *ctxA* counts.

Furthermore, by measuring the relative abundance of toxigenic and O1 serogroup strains in the total *V. cholerae* population as well as *V. metoecus* abundance, this assay provides information about intraspecies and interspecies population dynamics, which could yield insights into variations in the natural abundance of toxigenic strains. This assay has both a low detection limit (three copies per reaction) and higher specificity than comparable assays. Besides being the first assay to allow molecular quantification of *V. metoecus*, it is also the first one to have been extensively tested environmental samples for *V. cholerae*, by measuring the absolute abundance of the species and its O1 serogroup and toxigenic subpopulations in areas endemic for cholera and where the bacterium is undetectable by culture-based techniques over the course of several months.

## MATERIALS AND METHODS

### Bacterial cultures and DNA template preparation

Different isolates of *Vibrio* spp that were used to validate primer specificity and assay sensitivity in this study are listed in TABLE 1 and TABLE S1. All *Vibrio* strains were grown in tryptic soy broth (TSB) (Becton Dickinson) with 1.0% NaCl (BDH), incubated at 30 °C and 200 rpm. For non-*Vibrio* strains (TABLE S1) TSB without 1.0% NaCl was used under the same growth conditions. Genomic DNA was extracted from overnight cultures using the DNeasy Blood and Tissue Kit (QIAGEN) and quantified using the Quant-iT PicoGreen dsDNA Assay Kit (Molecular Probes) with a Synergy H1 microplate reader (BioTek).

**TABLE 1.** Bacterial strains used to validate specificity of the primers and probes designed to detect *V. cholerae* and its toxigenic serogroup O1 and *V. metoecus*.

| Species | No. of strains | Target genes |  |  |  |
| --- | --- | --- | --- | --- | --- |
|  |  | *viuB* | *ctxA* | *rfbO1* | *mcp* |
| *Vibrio cholerae* non O1 | 20 | + | - | - | - |
| *Vibrio cholerae* O1 | 8 | + | + | + | - |
| *Vibrio parahaemolyticus* | 1 | - | - | - | - |
| *Vibrio vulnificus* | 3 | - | - | - | - |
| *Vibrio metoecus* | 10 | - | - | - | + |
| *Vibrio mimicus* | 3 | - | - | - | - |
| *Escherichia coli* | 3 | - | - | - | - |
| *Pseudomonas aeruginosa* | 3 | - | - | - | - |

+, indicates all strains (100%) were positive by qPCR
-, indicates no amplification

### DNA extraction from biomass and isolation of organisms from environmental water samples

Water samples were collected from Bangladesh (a cholera-endemic region) and the USA East Coast (a cholera-free region) (FIG 1) at different time points. Environmental water samples from Oyster Pond (Falmouth, MA, USA) were collected during the months of June to September, 2008 and 2009, as previously described (Kirchberger et al., 2016). Briefly, triplicate samples were obtained at a 0.5 m depth and a distance of 5 m from each other. Samples from 2009 were size fractionated, where ten litres of water were first filtered through a 63 μm nylon mesh net to capture large particles such as zooplankton. Large particles were crushed in a 50 ml tissue grinder after transfer using 20 ml of sterile filtered local water. Two milliliters of crushed material (equivalent to one litre of water) was resuspended in 48 ml of sterile filtered local water and pushed through a 4.5 cm Millipore Durapore filter (0.22 μm pore size) using a polypropylene syringe.

**FIG 1.**
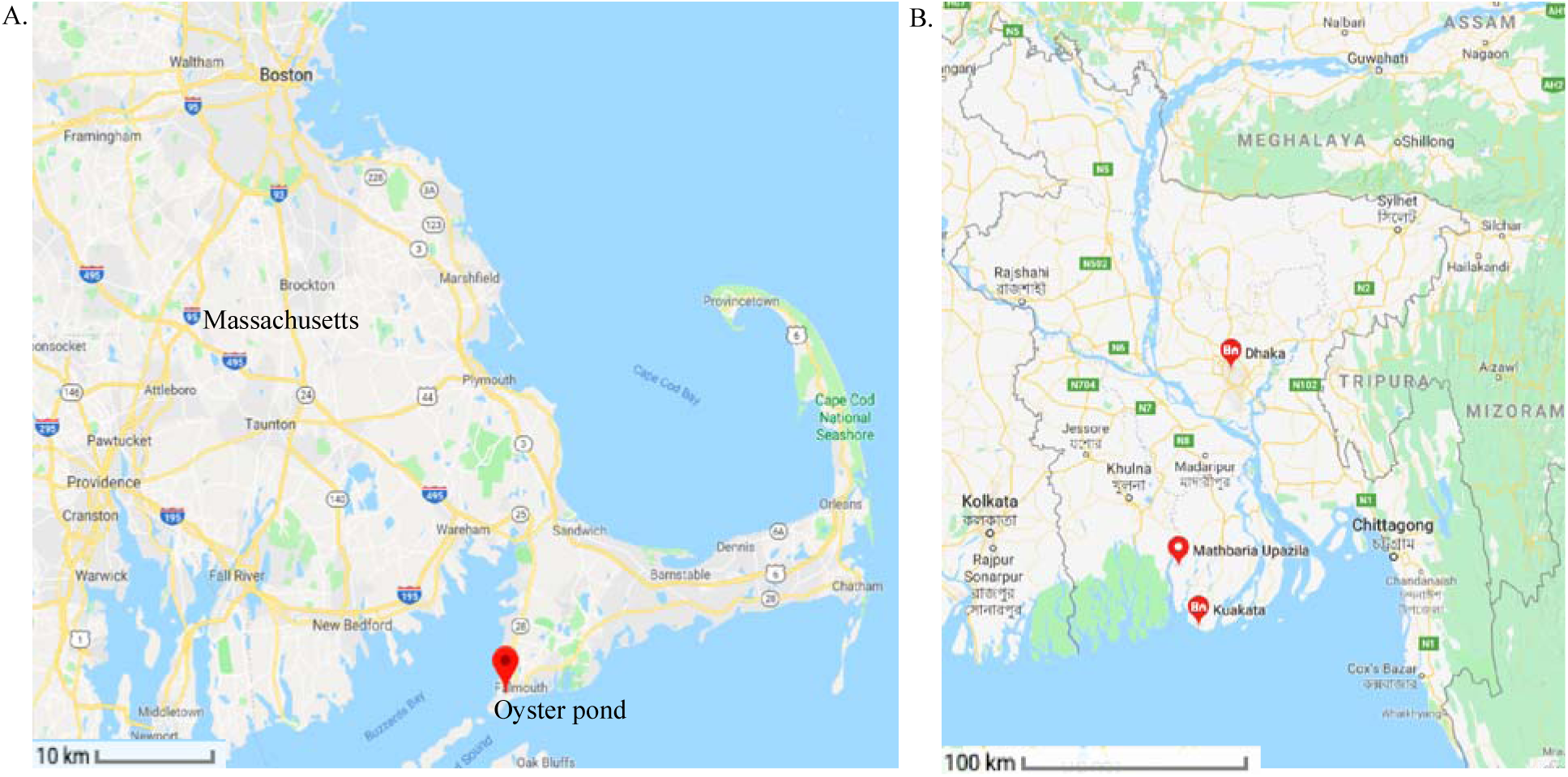
Sampling sites for environmental water samples collected to evaluate the qPCR assay developed in this study. A) Oyster Pond (Falmouth, Massachusetts, USA) a non-endemic site for cholera. B) Map of Bangladesh, identifying the two coastal regions (Kuakata and Mathbaria) and inland region (Dhaka) where samples were collected.

Similarly, one litre of water passed through the mesh net was pushed through a series of in-line 4.5 cm Millipore Durapore filters (sizes 5 μm, 1 μm, and 0.22 μm) using a peristaltic pump. All disposable equipment was sterile, and all filter casing and tubing was sterilized before sampling. DNA extraction from the filters using a QIAGEN DNeasy Blood and Tissue Kit was performed as follows: 0.25 g of sterile zirconium beads were added to cut-up filter pieces in a 1.5 ml screw cap tube with 360 μl Cell Lysis Buffer ATL, and bead beating performed for 30 sec at maximum speed. Proteinase K (40 μl) was added, and the tubes were vortexed for several seconds. Further steps followed the instructions of the manufacturer. Environmental strains of *V. cholerae* and *V. metoecus* were isolated from the August and September 2009 filters, as previously described (Kirchberger et al., 2016).

Three different regions in Bangladesh were selected to collect environmental water samples: two coastal regions (Kuakata and Mathbaria) and an inland region (Dhaka). Collection of water samples and extraction of DNA from filters from coastal Bangladesh sites (Kuakata and Mathbaria) were done at a single time point (May 2014) using the same protocol as the Oyster Pond sampling (Kirchberger et al., 2016). The Dhaka samples were collected from Rampura, Dhaka, Bangladesh (FIG 1) bi-weekly from October 2015 to March 2016. Water samples (50 ml) were collected using 60 ml sterile polypropylene syringe and filtered through 0.22 μm Sterivex filters (Millipore). Total DNA extraction from the biomass on the filters was done by following four consecutive steps: cell lysis and digestion, DNA extraction and DNA concentrating, and washing according to the protocol described by Wright *et al* (Wright et al., 2009). Environmental strains of *V. cholerae* were isolated from the water samples collected in Dhaka by following the protocol previously described (Huq et al., 1990).

To reduce impurities that can act as inhibitors during PCR amplification, all extracted DNA samples were further treated with One step PCR inhibitory removal kit (The Epigenetics Company, ZYMO Research), following the protocol in the user manual. Treated samples were kept at −20° C for further analysis.

### Design and evaluation of primers and fluorogenic probes for real–time qPCR

To design a primer set suitable for the amplification of products that are unique to *V. cholerae* strains, protein-coding genes from a dataset of 77 *Vibrio* genomes were analyzed. Initial screening for genes which are not shared between *V. cholerae* and *V. metoecus* as well as their close relatives was performed on a dataset of consisting of the genomes of 42 *V. cholerae*, 10 *V. metoecus* and 25 other *Vibrio* strains. OrthoMCL (Li et al., 2003) was used to cluster all protein-coding genes into gene families based on 30% amino acid identity (Rost, 1999), resulting in sets of genes found exclusively in *V. cholerae* or *V. metoecus* and not shared with other species, respectively. After removal of multi-copy genes, remaining genes were aligned using CLUSTALW 2.0 (Larkin et al., 2007) and manually inspected for sites appropriate for the design of qPCR probes. Optimal sites for non-degenerate primer/probe design were found in the *viuB* gene encoding the vibriobactin utilization protein B (NP_23184.1) for *V. cholerae* (Butterton and Calderwood, 1994; Wyckoff et al., 2007). Primers (forward and reverse) and probe (TABLE 2) for a 77-bp product were designed using the software tool PrimerQuest from integrated DNA technologies (IDT, Iowa, USA) according to supplied guidelines.

**TABLE 2.**
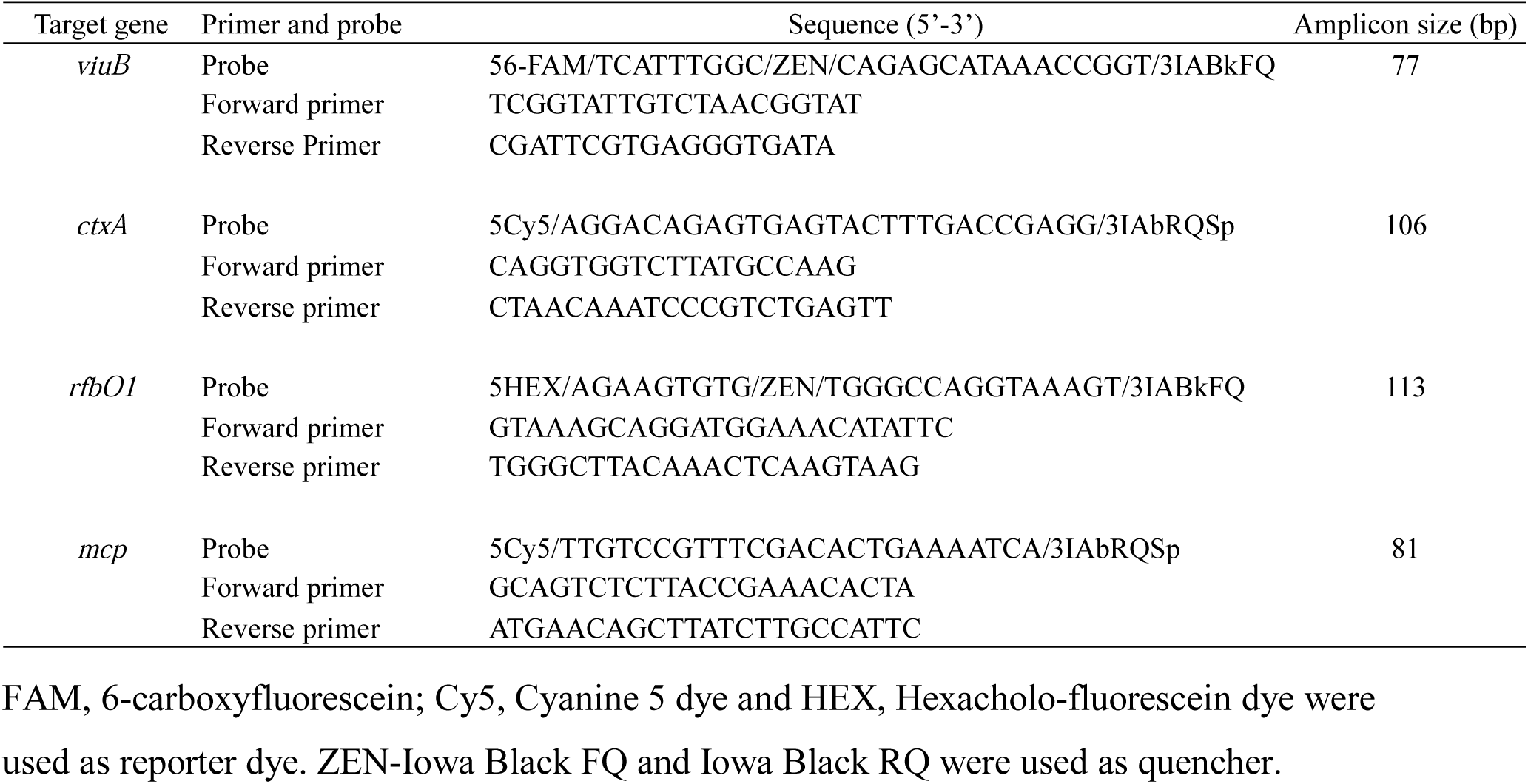
Target genes and sequences of primers and probes used in this study.

To design primers specific for *V. metoecus*, Intella (https://www.vound-software.com/) was used to analyze the unique gene contents of *V. metoecus*. Alignments of the sequence of alleles from single copy gene families present in all *V. metoecus* in the dataset were performed by using an in-house script (available upon request). The gene encoding a methyl-accepting chemotaxis protein (MCP) (EE X66169.1) was selected for the presence of ideal primer and probe sites (Integrated DNA technologies), as designed using the software PrimerQuest tool from Integrated DNA technologies (IDT, Iowa, USA). This gene was targeted using the probe 5’-/5Cy5/TTG TCC GTT TCG ACA CTG AAA TCA/3IAbRQSp/-3’, and forward and reverse primers, 5’-GCA GTC TCT TAC CGA AAC ACT A-3’ and 5’-ATG AAC AGC TTA TCT TGC CAT TC-3’, respectively, yielding an 81-bp product. The designed primers were tested for specificity by end point PCR with spiked water samples.

For estimation of toxigenic *V. cholerae* abundance, 106 bp of the *ctxA* gene (TABLE 2) was targeted, as part of the genetic element encoding the major virulence factor cholera toxin and thus one of the signature genes for toxigenic potential in *V. cholerae* (Mekalanos et al., 1983). In the case of the *V. cholerae* O1 serogroup, the target was a 113 bp product of the *rfbO1* gene (TABLE 2) as it detects explicitly *V. cholerae* belonging to the O1 serogroup, which includes the vast majority of strains responsible for past and ongoing cholera pandemics. Primers and probes for both of these genes were designed in this study to ensure compatibility of the assay.

### Real-time qPCR amplification

Dynamite qPCR Mastermix used in this study is a proprietary mix, developed and distributed by the Molecular Biology Service Unit at the University of Alberta, Edmonton, AB, Canada. It contains Tris (pH 8.3), KCl, MgCl_2_, glycerol, Tween 20, DMSO, dNTPs, ROX as a normalizing dye, and antibody inhibited *Taq* polymerase. The volume of the PCR reaction was 10 μl containing 5 μl of 2× Dynamite qPCR master mix, 1 μl of each of 500 nM primer-250 nM probe mix, 1 μl of molecular grade water and 2 μl of DNA template. Real-time quantitative PCR was performed under the following conditions: initial primer activation at 95 °C for 2 min followed by 40 cycles of 95 °C for 15 s, 60 °C for 1 min in the Illumina Eco Real-Time PCR system. The assay includes standards of known copy number and negative control with no template added to assess the potential presence of contamination (FIG 2).

**FIG 2.**
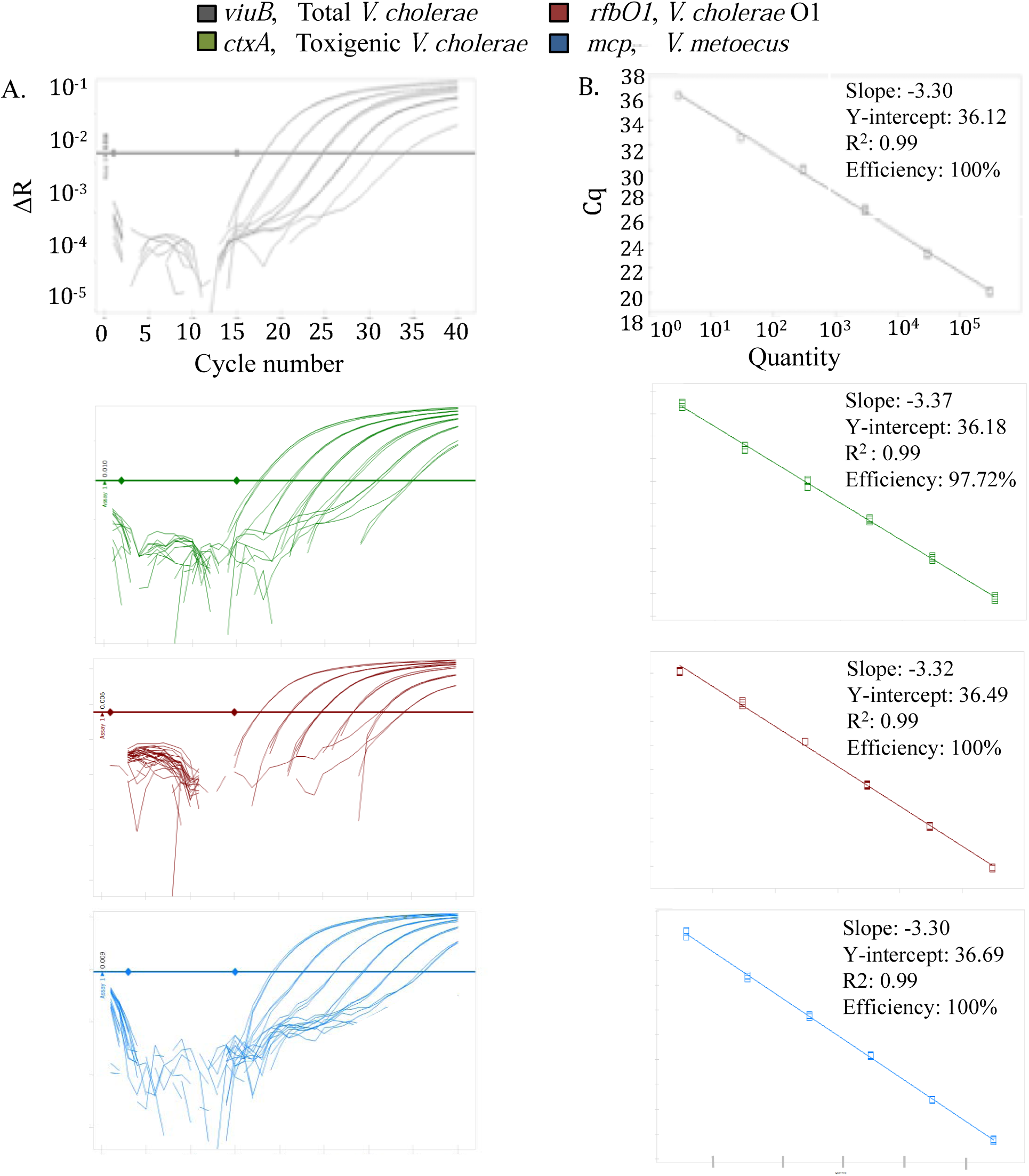
Multiplex real-time qPCR for simultaneous detection and quantification of *V. cholerae* and *V. metoecus*. Four gene markers with fluorogenic probes were used: *viuB* (*V. cholerae* specific), *ctxA* (toxigenic *V. cholerae* specific), *rfbO1* (*V. cholerae* O1 specific) and *mcp* (*V. metoecus* specific). Template DNA was purified from reference cultures (*V. cholerae* N16961 and *V. metoecus* RC341) and was serially diluted in 10-fold increments to yield

### Generation of standard curves and calculation of qPCR efficiency

Standard curves were prepared by amplifying gene sequences from corresponding reference strains for each target. For the preparation of standard curves, the *viuB, ctxA* and *rfbO1* genes of the pandemic *V. cholerae* El Tor O1 N16961 reference strain were used for total *V. cholerae* count, toxigenic *V. cholerae* and *V. cholerae* O1, respectively. To make the standard curve for the *V. metoecus* specific gene (*mcp*), the *V. metoecus* RC341 was used. Both strains were grown on LB agar (BD Difco, USA) with 0.5% NaCl at 30 °C for overnight and DNA extraction was done by DNeasy Blood and Tissue Kit (QIAGEN). Specific forward and reverse primers targeting each gene of interest were used for PCR amplification (TABLE 2). A standard PCR protocol was followed for this amplification: 1 μl each of 10 pmol forward and reverse primer, 0.4 μl of 10 mM dNTP-Mix (ThermoFisher), 0.4 μl Phire Hot Start II DNA Polymerase (ThermoFisher), 4 μl of 5× Phire Buffer, 12.2 μl of molecular biology grade water and 2 μl of template DNA. The PCR reaction was performed as follows: initial denaturation at 98 °C for 30 sec, followed by annealing at 55 °C for 5 sec and extension 72 °C for 1 min for 35 cycles and a final extension of 72 °C for 1 min. PCR products were purified using the Wizard SV Gel and PCR Clean-up System (Promega). The concentrations of amplified PCR products were measured using the Quant-iT PicoGreen dsDNA Assay Kit (Molecular Probes) and the Synergy H1 microplate reader (BioTek).

Calculations mentioned in the Applied Biosystems Guideline (Applied Biosystems) for creating qPCR standard curves were used for determining the mass of amplified gene templates that correspond to copy numbers of target nucleic acid sequences. A series of standards were prepared in which a gene of interest is present at 3×10^5^ copies, 3×10^4^ copies, 3×10^3^ copies, 3×10^2^ copies 30 and 3 copies per 2 μl of the template. Once prepared, the standards were stored in 100 μl aliquots at −80° C. The standard curve was generated by plotting the log value of calculated gene copies per reaction over the quantitative cycle value (Cq) (FIG 2). The Cq is described as the cycle at which the fluorescence from amplification exceeds the background fluorescence in the MIQE guideline (Bustin et al., 2009).

If a sample contains more targets, the fluorescence will be detected in earlier cycles; low Cq values represent higher initial starting copies of the target gene. The qPCR efficiency of the assay was calculated (FIG 2) using the following formula: Efficiency = 10^[-1/Slope]^ (http://efficiency.gene-quantification.info/) by Illumina Eco Real-Time PCR system software. concentration ranging from 3×10^6^ to 3 copies per reaction (from left to right). Fluorescence was measured in relative units. The panel (A) illustrates amplification curves and the panel (B) shows their corresponding standard curves. Each reaction was done in triplicate.

### Limit of detection (LOD) and impact of inhibition testing

The LOD of the assay was determined for each of the marker genes based on the standard curve of amplified genes from reference strains (*V. cholerae* N16961 and *V. metoecus* RC341) (FIG 2). The LOD of sample (per liter of water) before filtration was calculated from the LOD of the qPCR assay. We did not evaluate the quantifiable lowest minimum number of copies in the environmental water sample before any filtration step.

To test for qPCR inhibition, we compared the Cq values for 10× dilution of treated (with One step PCR inhibitory removal kit) extracted DNA samples from study sites and spiked positive and negative samples. The difference in Cq values between diluted samples were recorded (FIG S2).

### Specificity testing

The specificity of the qPCR assay was evaluated using genomic DNA from bacterial strains listed in TABLE 2. Non-O1 *V. cholerae* (20) from environmental sources, *ctxA* positive *V. cholerae* O1(8) from both clinical and environmental sources, and *V. metoecus* (10) from environmental sources were tested (TABLE S1). Three other *Vibrio* species, *V. parahaemolyticus, V. vulnificus* and *V. mimicus*, were also tested, as well as two non-*Vibrio* gammaproteobacteria, *Pseudomonas aeruginosa* and *Escherichia coli*.

## RESULTS AND DISCUSSION

### A specific and efficient multiplex qPCR assay to detect *V. cholerae* and *V. metoecus*

We developed a specific and sensitive qPCR method for the quantification of total *V. cholerae* and *V. metoecus*, as well as toxigenic and O1 serogroup *V. cholerae*, in their natural aquatic environments. Primers and probes developed for all four marker genes displayed 100% specificity (TABLE 1) using 51 bacterial strains, including 20 *V. cholerae* strains of various serogroups and toxigenic potential, as well as 10 *V. metoecus* and closely related *Vibrio* species. This assay was more specific than any previously published, with no cross-amplification between *V. cholerae* and other bacteria, including its closest relative, *V. metoecus*. The efficiency of each assay was 95% to 100% (R^2^ value 0.99 and slope −3.3 ± 0.07) (FIG 2).

The *viuB* and *mcp* markers are specific to *V. cholerae* and *V. metoecus* and present in single copies, facilitating quantification of the absolute abundance of these two species. Additionally, the *ctxA* and *rfbO1* markers are specific for detection of toxigenic and O1 serogroup strains, respectively. Previous qPCR studies often took advantage of these two genes but did not combine them, resulting in missed information required to correlate toxigenic *V. cholerae* and strains representing the PG lineage (Blackstone et al., 2007; Bliem et al., 2015). Although the region selected for *ctxA* gene amplification to indicate the presence of toxigenic *V. cholerae* is specific to this particular species (Goel et al., 2007), the presence of *ctxAB* has been reported in CTXΦ phages present in aquatic environment, which can be the source of *ctxA* positive results (Faruque et al., 1998). Multiple copies of *ctxA* can also be present in certain *V. cholerae* genomes, biasing quantification results (Faruque et al., 1994; Mekalanos, 1983). The simultaneous amplification of the single copy *rfbO1* specific for *V. cholerae* O1 strains compensates for these flaws, as the vast majority of cholera cases are caused by CTX positive strains of the O1 serogroup, and the co-occurrence of these two markers at similar levels allows accurate quantification of toxigenic O1 strains.

There are many rapid assays already being used to detect certain serotypes of *V. cholerae*, but most of them are confined to qualitative detection and do not provide any data on the abundance of this organism in the environment (Chua et al., 2011; Goel et al., 2007; Koskela et al., 2009; Singh et al., 2002). A few available assays are quantitative, but the limitations in sensitivity are >5 to 50 gene copies per reaction (Rashid et al., 2017b; Vezzulli et al., 2015), and they detect no more than two marker genes, or lack specificity (Gubala, 2006). For example, a real-time PCR assay using four different genetic markers i.e. *rtxA, epsM, ompW* and *tcpA* to detect *V. cholerae* also detected *V. mimicus* non-specifically by amplification of *ompW* (Gubala, 2006). Other genes currently used for detection and/or quantification of *V. cholerae* such as *hlyA, zot, ompU, toxR, groEl*, also suffer from a lack of species specificity (Fykse et al., 2007; Goel et al., 2007; Singh et al., 2002). Many of these genes share sequence similarity with homologs in closely related species or can be present in multiple copies in *Vibrio* genomes, and therefore can pose problems for specific detection and quantification. Moreover, there is no published assay that detects and quantifies the abundance of *V. metoecus* along with *V. cholerae.* Most assays have been evaluated by artificially spiking water samples with known concentrations of laboratory strains, and none has been directly applied to a significant number of environmental samples in a region endemic for cholera.

It is important for any environmental application to determine the sample limit of detection. This value represents the lowest quantity of the target DNA that can be reliably detected and quantified in a certain volume of sample at a probability level of 95% (Bustin et al., 2009). In this study, we found the analytical detection limit for all four gene markers to be three copies per reaction (FIG 2), with 1.5×10^6^ copies/L of water as the lowest detectable number without filtration, which is comparable to previous studies (Fykse et al., 2007). Concentrating samples by filtration of large volumes of water (0.05 L to 10 L, based on sampling at four sites in this study) made it possible to lower the detection limit to 9.0×10^2^ copies/L for Oyster Pond (USA) samples after filtering 566 ml of water (FIG 3 and FIG 4B). After filtration, lowest detection limit of 3.0×10^3^ copies/L (from 10 L of water) (FIG 5 and FIG 6A) and 1.5×10^4^ copies/L (from 50 ml of water) (FIG 7) were determined for samples from the Kuakata and Dhaka (Bangladesh) sites, respectively.

**FIG 3.**
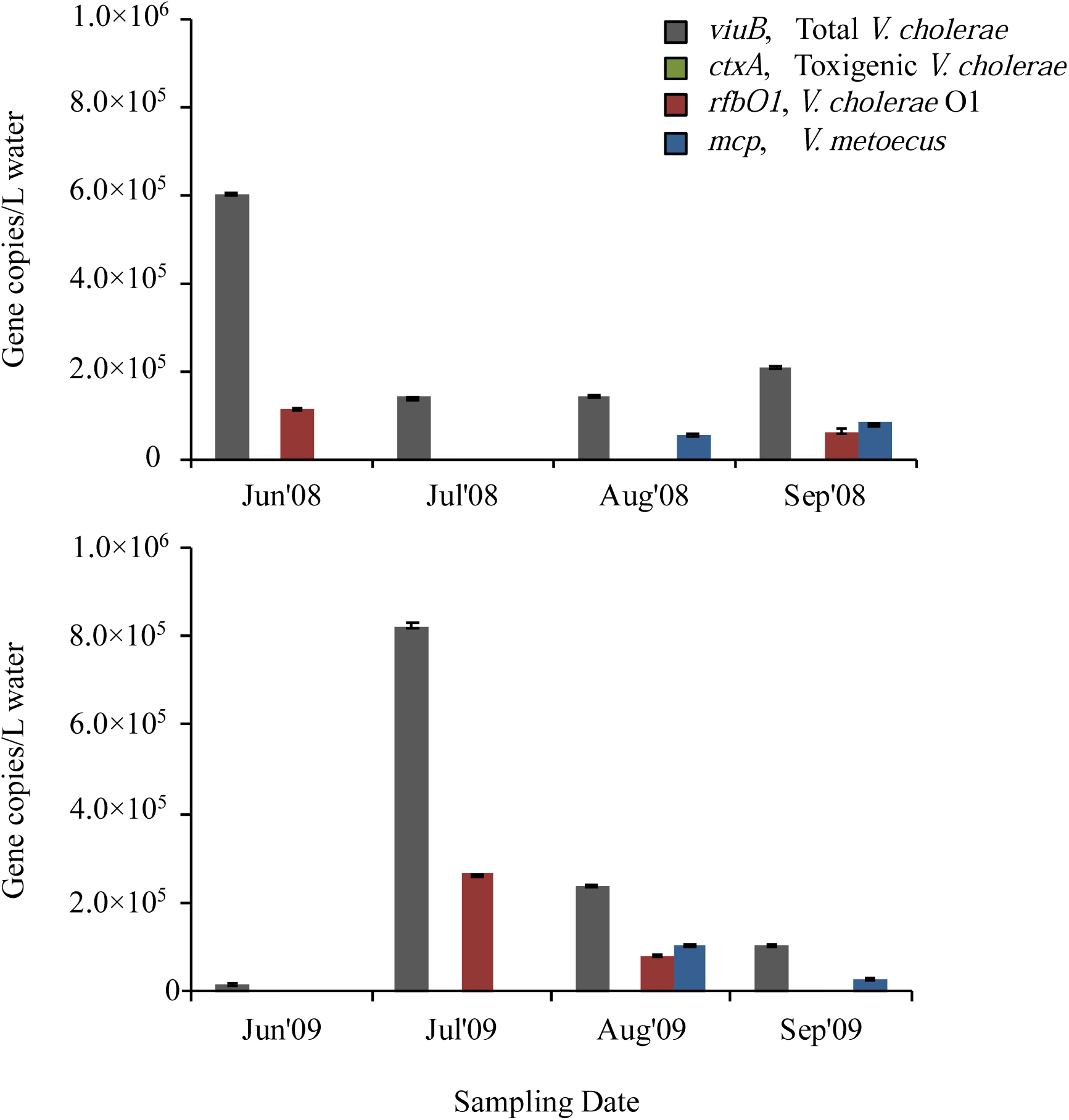
Temporal variation of abundance for *V. cholerae* along with its toxigenic and serogroup O1 subpopulations and its close relative *V. metoecus* in Oyster Pond, MA, USA. Environmental water samples collected during the months of June to August in two successive years, 2008 and 2009, were analyzed using the developed qPCR assay. The *viuB* gene was used to quantify total *V. cholerae*; *ctxA* and *rfbO1* were used to measure toxigenic *V. cholerae* and *V. cholerae* O1, respectively; the abundance of *V. metoecus* was estimated using the *mcp* gene. All four genes were tested for each sample and the absence of a bar in the graph denotes that the target gene was absent or present below the detection limit of the assay. Each qPCR reaction was run in triplicate. Mean values are shown with error bars indicating the standard deviation between three technical replicates. *ctxA* could not be detected at any time sampled at this site.

**FIG 4.**
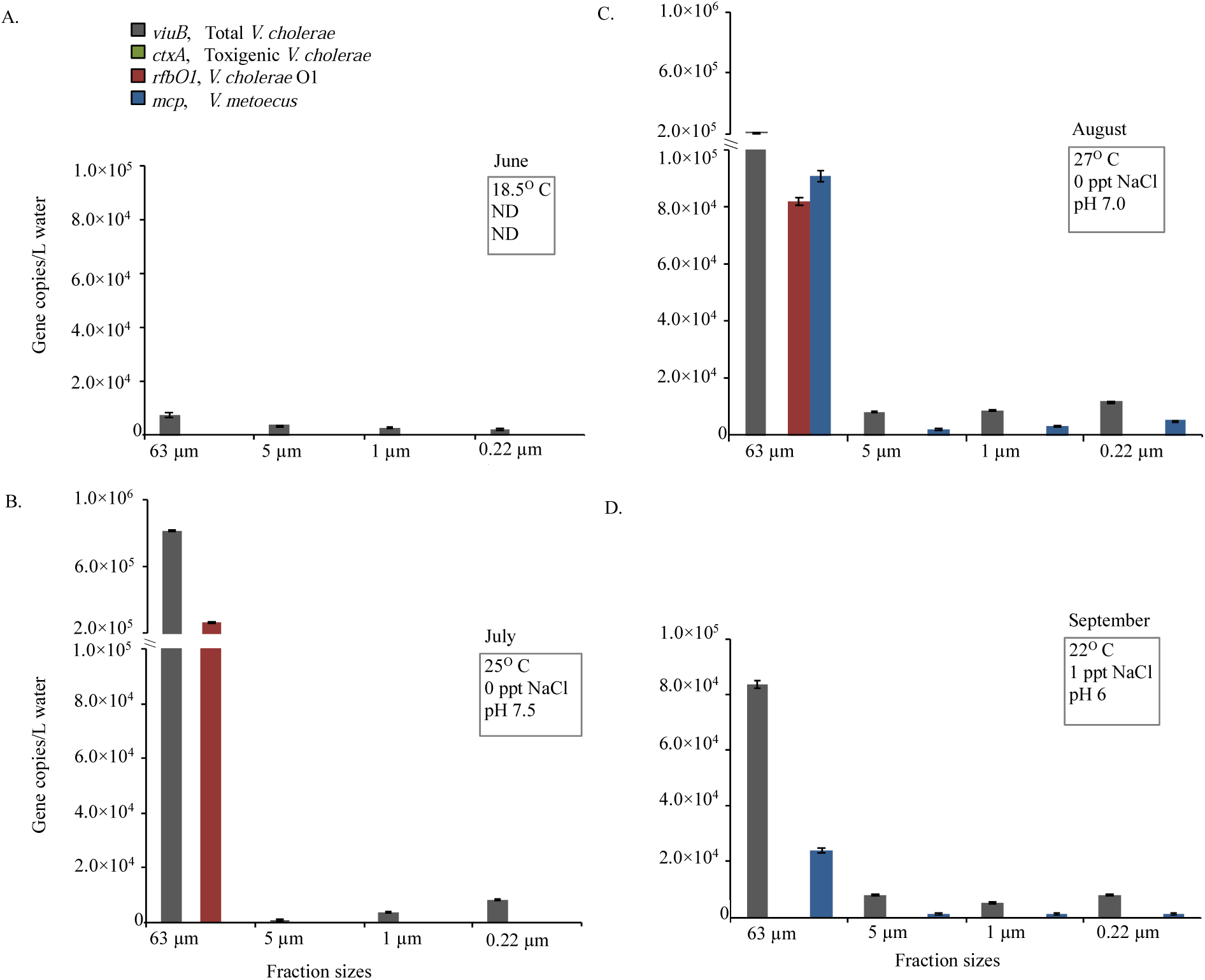
Distribution in different water fraction sizes of *V. cholerae* along with its toxigenic and serogroup O1 subpopulations and its close relative *V. metoecus* in Oyster Pond, MA, USA. Environmental water samples were collected during the months of June (A), July (B) August (C) and September (D) in 2009. These samples were fractionated by size through sequential filtration and bacteria were quantified by qPCR of marker genes on DNA extracted from the filters. The *viuB* gene was used to quantify total *V. cholerae*; *ctxA* and *rfbO1* were used to measure toxigenic *V. cholerae* and *V. cholerae* O1, respectively; the abundance of *V. metoecus* was quantified using the *mcp* gene. Each qPCR reaction was run in triplicate. Mean values are shown with error bars indicating the standard deviation between technical replicates. Temperature, pH and salinity of water collected each month are shown in boxes on the upper right corner of each graph. ND indicates not done.

**FIG 5.**
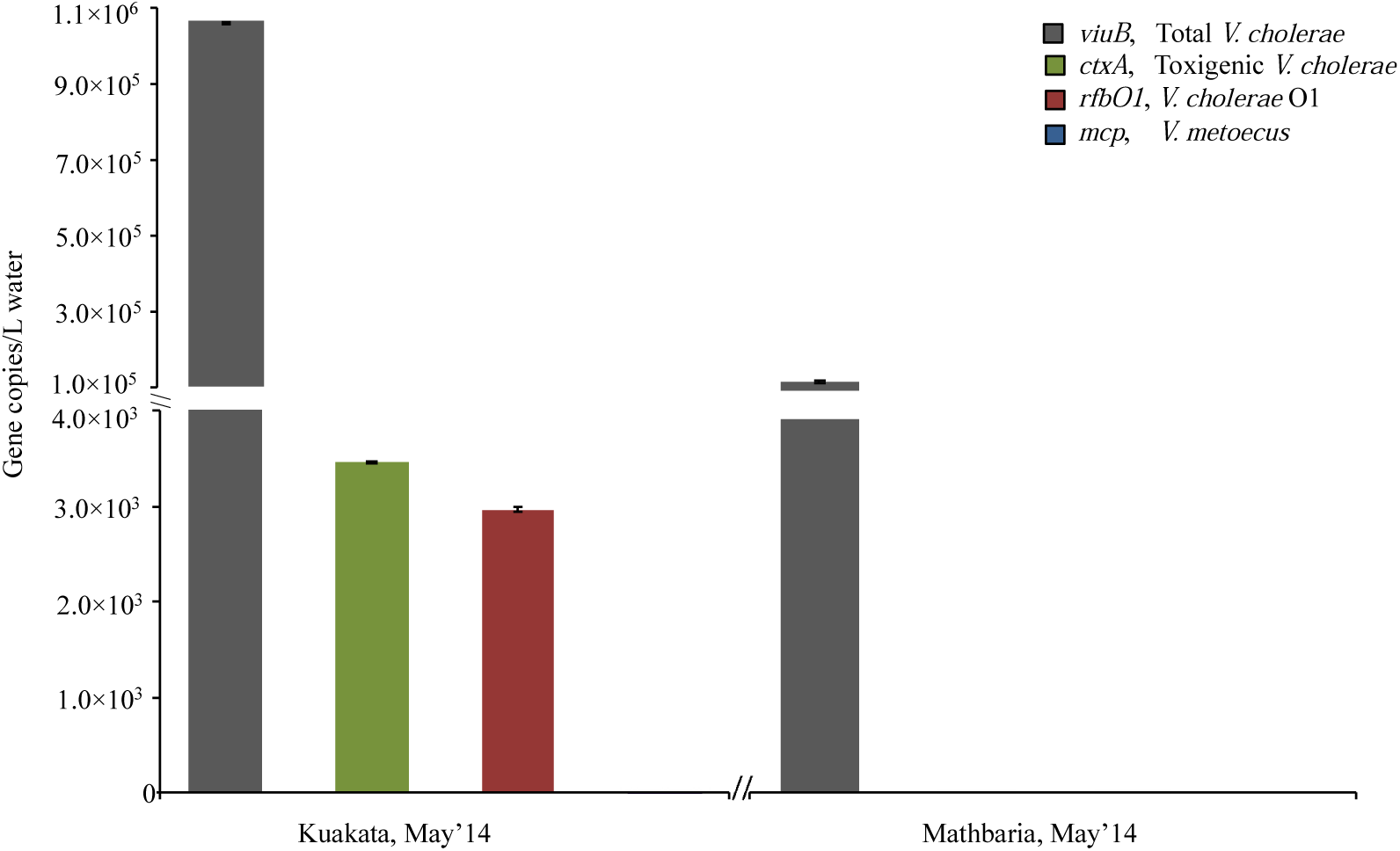
Abundance of *V. cholerae* along with its toxigenic and serogroup O1 subpopulations and its close relative *V. metoecus* in two different coastal regions in Bangladesh. Environmental water samples were collected from Kuakata and Mathbaria during the month of May in 2014 and bacteria were quantified by qPCR of marker genes. The *viuB* gene was used to quantify total *V. cholerae*; *ctxA* and *rfbO1* were used to measure toxigenic *V. cholerae* and *V. cholerae* O1, respectively; the abundance of *V. metoecus* was quantified using the *mcp* gene. Each qPCR reaction was run in triplicate. Mean values are shown with error bars indicating the standard deviation between technical replicates.

**FIG 6.**
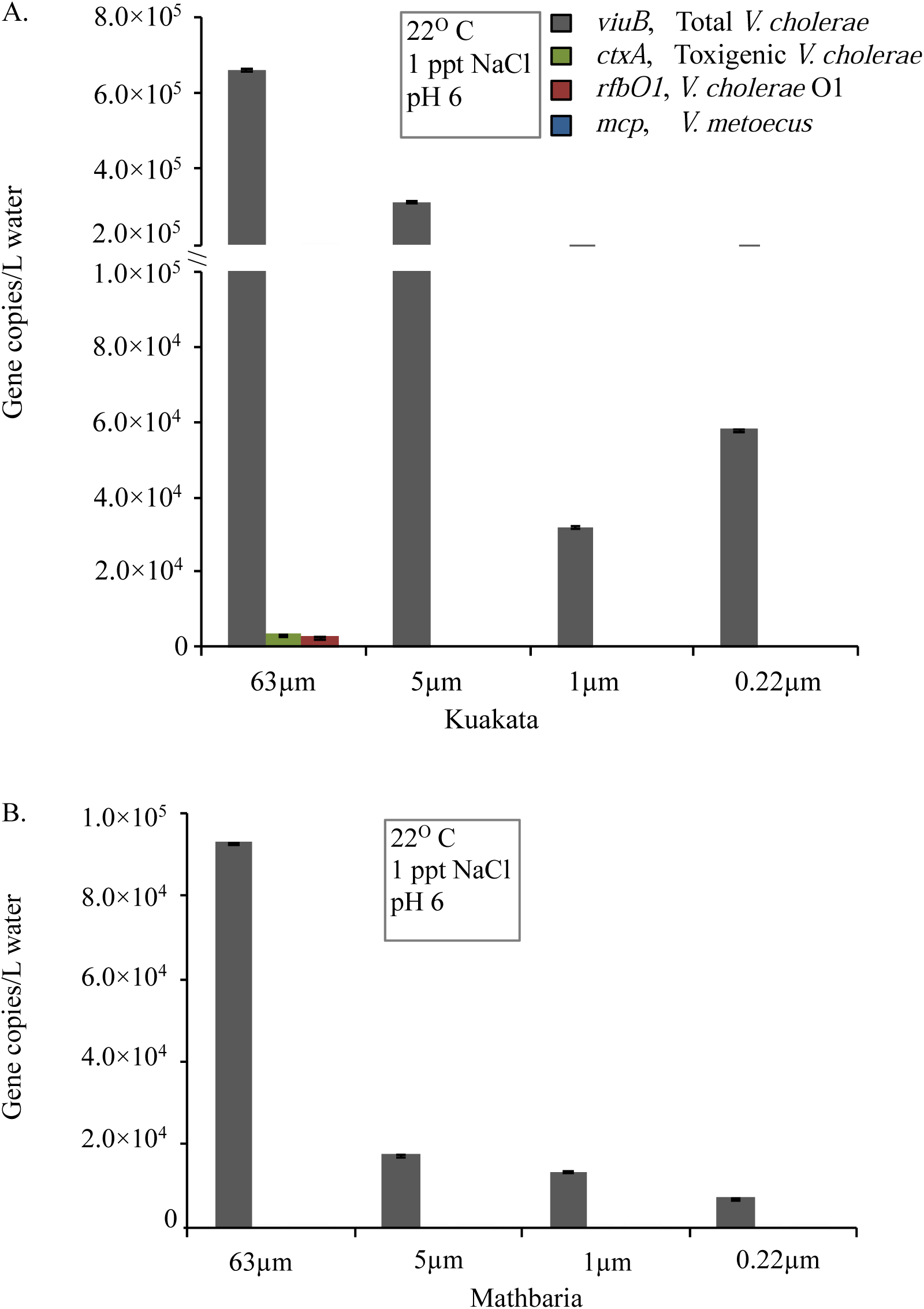
Distribution in different water fraction sizes of *V. cholerae* along with its toxigenic and serogroup O1 subpopulations and its close relative *V. metoecus* in two different coastal regions in Bangladesh. Environmental water samples were collected from A) Kuakata and B) Mathbaria during the month of May in 2014 and bacteria were quantified by qPCR of marker genes. The *viuB* gene was used to quantify total *V. cholerae*; *ctxA* and *rfbO1* were used to measure toxigenic *V. cholerae* and *V. cholerae* O1, respectively; the abundance of *V. metoecus* was quantified using the *mcp* gene. Each qPCR reaction was run in triplicate. Mean values are shown with error bars indicating the standard deviation between technical replicates.

**FIG 7.**
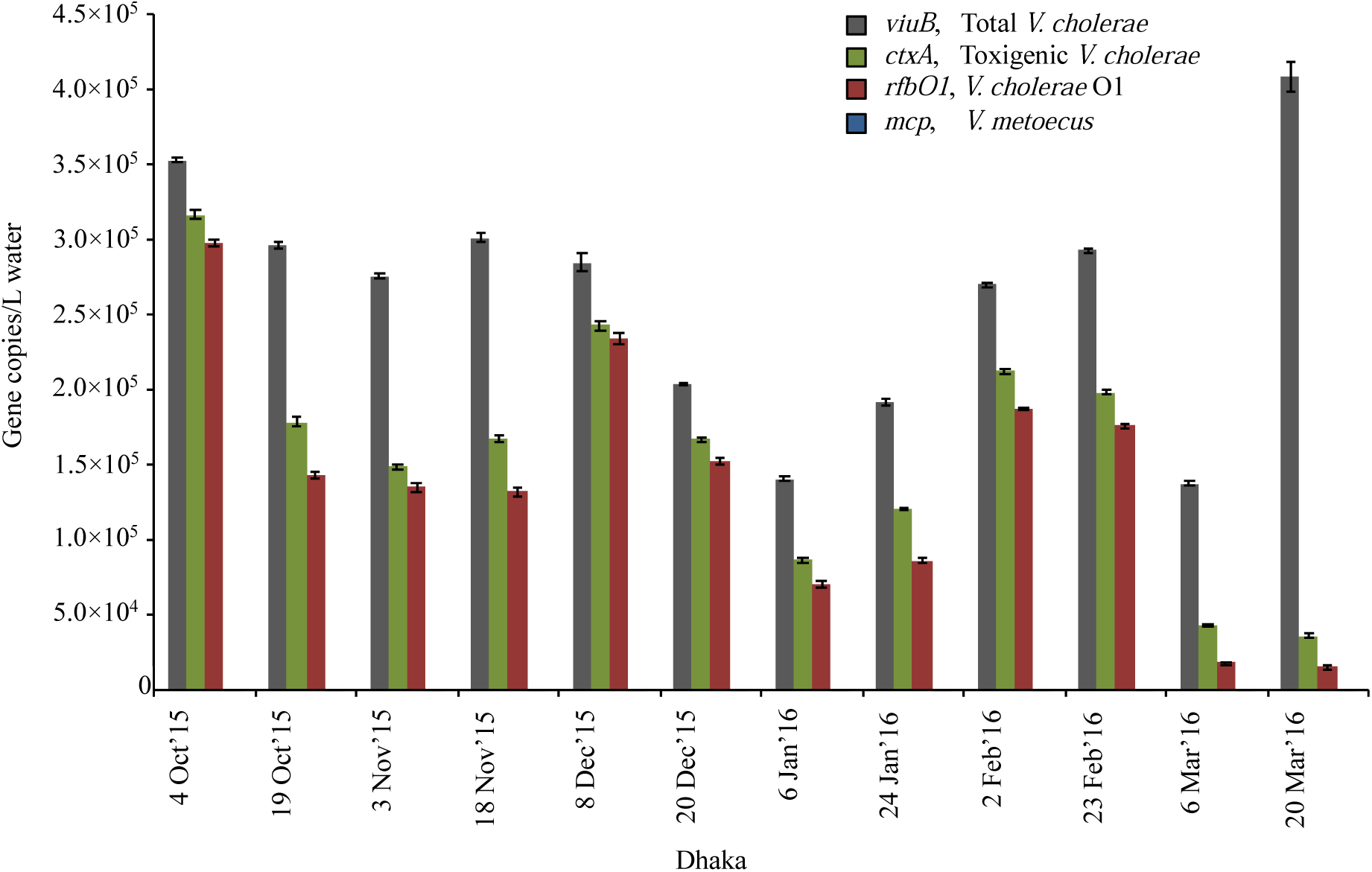
Temporal variation of abundance for *V. cholerae* along with its toxigenic and serogroup O1 subpopulations and its close relative *V. metoecus* in a central region of Bangladesh (Dhaka). Environmental water samples were collected bi-weekly from the months of October 2015 to March 2016 and bacteria were quantified by qPCR of marker genes. The *viuB* gene was used to quantify total *V. cholerae*; *ctxA* and *rfbO1* were used to measure toxigenic *V. cholerae* and *V. cholerae* O1, respectively; the abundance of *V. metoecus* was quantified using the *mcp* gene. Each qPCR reaction was run in triplicate. Mean values are shown with error bars indicating the standard deviation between technical replicates.

The sensitivity of PCR-based assays for quantifying *V. cholerae* from environmental samples is highly dependent on efficient DNA extraction and removal of potential inhibitors. All the environmental DNA samples were treated with One step PCR inhibitory removal kit after DNA extraction to reduce residual inhibitors. Samples were assessed in ten-fold dilutions (1/10) to determine the effect of inhibitors in the assay. The 10× dilution of samples shifted the Cq values by 3.3 cycles ± 0.05 (FIG S2.), consistent with no significant inhibition.

### *V. cholerae* and *V. metoecus* co-occur seasonally in a temperate coastal location

With an optimized multiplex qPCR assay, it was possible for the first time to determine the abundance of *V. cholerae* and *V. metoecus* simultaneously in water reservoirs in cholera non-endemic areas (USA). *V. cholerae* is an ubiquitous member of bacterial communities in temperate and tropical aquatic environments around the world (Kaper et al., 1995) whereas *V. metoecus* is a recently described species, which has been co-isolated with *V. cholerae* from the USA East Coast aquatic environment and associated with eel fish in Spain, as well as clinical cases from around the USA (Boucher et al., 2011; Carda-Diéguez et al., 2017; Kirchberger et al., 2016; Kirchberger et al., 2014).

To compare the qPCR assay to culture-dependent methods of detection, it was applied to samples from which *V. cholerae* and *V. metoecus* had been isolated by cultivation and samples from which no organisms had been isolated (TABLE 3). These water samples were collected in summer (June to September) over two successive years (2008 and 2009) from Oyster Pond in Falmouth (MA) on the East Coast of the USA. Using our novel multiplex assay on DNA extracted from biomass of the same samples, we found the abundance of *V. cholerae* was 1.4×10^5^ to 8.2×10^5^ copies/L in Oyster Pond from June to September in 2008 and 2009 (FIG 3). Strikingly, *V. metoecus* was only detected at the end of the season (the months of August and September) in both years, at abundances of approximately 5.6×10^4^ to 1.0×10^5^ copies/L (FIG 3). *V. cholerae*, on the other hand, was consistently more abundant than *V. metoecus* and present throughout the summer. Using a culture-based approach, it has been suggested that *V. cholerae* is ten times more abundant than *V. metoecus* at that particular location (Kirchberger et al., 2016). Based on the qPCR approach used here, *V. cholerae* was approximately three times more abundant than *V. metoecus*, suggesting that the former is more readily culturable than the latter (FIG 3). Moreover, *V. metoecus* was neither isolated by conventional culture method nor detected by qPCR in Bangladesh, suggesting a different geographical distribution than *V. cholerae* (FIG 5).

**TABLE 3.** Comparative result of conventional culture method and developed qPCR technique used in this study.

| Site | Time | Cultivation |  |  | qPCR |  |  |
| --- | --- | --- | --- | --- | --- | --- | --- |
|  |  | *V. cholerae* | *V. cholerae* O1 | *V. metoecus* | *V. cholerae* | *V. cholerae* O1 | *V. metoecus* |
| Kuakata | May'14 | + | - | - | + | + | - |
| Mathbaria | May'14 | + | - | - | + | - | - |
| Dhaka | Oct'15 | + | + | - | + | + | - |
|  | Nov'15 | + | - | - | + | + | - |
|  | Dec'15 | + | - | - | + | + | - |
|  | Jan'16 | + | - | - | + | + | - |
|  | Feb'16 | + | - | - | + | + | - |
|  | Mar'16 | + | - | - | + | + | - |
| Oyster Pond | Jun'08 | ND | ND | ND | + | - | - |
|  | Jul'08 | ND | ND | ND | + | + | - |
|  | Aug'08 | ND | ND | ND | + | + | + |
|  | Sep'08 | ND | ND | ND | + | - | + |
| Oyster Pond | Jun'09 | ND | ND | ND | + | - | - |
|  | Jul'09 | ND | ND | ND | + | + | - |
|  | Aug'09 | + | - | + | + | + | + |
|  | Sep'09 | + | - | + | + | - | + |

+, indicates samples were positive for corresponding organism
-, indicates there was no isolation and/or negative by qPCR
ND, indicates not done

### O1 serogroup strains are important members of a temperate coastal *V. cholerae* population

The presence/absence of toxigenic or O1 serogroup *V. cholerae* was determined using two sets of primers targeting *ctxA* and *rfbO1*. There was no amplification of *ctxA* in the Oyster Pond samples, but *rfbO1* was present sporadically: 1.1×10^5^ copies/L were present in June 2008 and 6.2×10^4^/L in September 2008 (FIG 3), while 2.6×10^5^ copies/L *rfbO1* were detected in July 2009 and 8.2×10^4^ copies/L in August of 2009 (FIG 3). The proportion of *rfbO1* positive *V. cholerae* was, on average, ∼18% of total *V. cholerae* in Oyster pond over these periods of sampling, ranging from 15% to 21% in individual samples. O1 serogroup of *V. cholerae* were consistently found during the four warmest months (June to September) of 2008 and 2009 (FIG 3). Interestingly, in water samples that were fractionated by size (2009), *rfbO1* was detected only in the largest size fraction (>63 μm) (FIG 4).

No O1 serogroup strains could be obtained from Oyster pond samples by conventional culture methods, despite the isolation of over 385 *V. cholerae* strains (Kirchberger et al., 2016) (TABLE 3). *V. cholerae* enters into VBNC form due to environmental stress condition, where cells remain alive but cannot be revived using standard cultivation methods (Colwell, 2000). During inter-epidemic periods, toxigenic *V. cholerae* are believed to be maintained in low numbers, attaching to particles in a state of VBNC (Sultana et al., 2018).

The inability to isolate *V. cholerae* O1 in most of the samples indicates that culture-based methods could lead to an underestimation of the occurrence of *V. cholerae* O1 in environmental reservoirs due to their low abundance or VBNC state (Bliem et al., 2015). The inability to isolate *V. cholerae* O1 strains in 2009 samples collected from Oyster pond together with their detection at a relatively high abundance (TABLE 3 and FIG 3) by qPCR likely indicates the presence of VBNC cells, questioning the assumption that *V. cholerae* O1 is rare in cholera-free regions (Islam et al., 2013).

### Toxigenic O1 serogroup strains are constantly present at dangerous levels in Dhaka freshwater

In Bangladesh, data on the incidence of cholera show that the disease occurs year round in the Ganges delta region of Bangladesh with seasonal peaks typically before (March to May) and after (September to December) monsoon (Alam et al., 2006; Faruque et al., 2005b). However, using culture-based methods, direct isolation of toxigenic *V. cholerae* O1 was not readily possible even during the peak season in aquatic reservoirs unless further enrichment in an alkaline peptone water medium (Huq et al., 1990). To determine if culture-based monitoring is misleading, bi-weekly sampling was done over six consecutive months in the Rampura area of Dhaka city in Bangladesh.

Biomass was collected from water samples by filtration on 0.22 μm membranes with no size fractionation. The abundance of the *viuB* marker gene, which corresponds to the total *V. cholerae* population, ranged from 1.3×10^5^ to 4.0×10^5^ copies/L from October 2015 to March 2016 (FIG 7). The *ctxA* gene used to track toxigenic *V. cholerae* was found at 3.6×10^4^ copies/L to 3.2×10^5^ copies/L and the *rfbO1* gene used to track *V. cholerae* O1 was detected at similar levels, ranging from 1.5×10^4^ copies/L to 2.9×10^5^ copies/L, corresponding to 50-60% of the total *V. cholerae* population (FIG 7). There was consistent occurrence of *V. cholerae* and its toxigenic and serogroup O1 subpopulations at similar abundances throughout the six months of sampling, suggesting that both genes were found in the same cells, which is typical of 7^th^ pandemic *V. cholerae* El tor strains currently responsible for most cholera cases in Bangladesh (Chun et al., 2009). Conventional culture methods only recovered *V. cholerae* O1 from October 2015, and *V. cholerae* non-O1 was isolated from each of these six months (TABLE 3).

Unexpectedly, analysis of water samples from the Rampura area (Dhaka, Bangladesh) (FIG 1) revealed persistent toxigenic *V. cholerae* O1 at a low abundance (1.3×10^5^ to 4.0×10^5^ copies/L), but a high proportion of the total *V. cholerae* population (up to 84%) (FIG 7). The infectious dose of *V. cholerae* in humans being in the order of 10^3^ to 10^8^ bacterial cells (Schmid-Hempel and Frank, 2007); the water in this region is a permanent reservoir of toxigenic *V. cholerae* that can readily cause potential outbreaks when ingested with contaminated water or food. Water bodies around Dhaka city are surrounded by a dense population which extensively interact with them, potentially resulting in the circulation of pathogenic *V. cholerae* in that particular environment, even outside of periods in which cholera cases are frequent. Usually, this megacity experiences two seasonal outbreaks of cholera before the monsoon in March to May and just after the monsoon in September to November (Alam et al., 2011). In this study we also observed higher counts of toxigenic *V. cholerae* in early October (2.9×10^5^ copies/L) and higher counts of total *V. cholerae* (> 4.0×10^5^ copies/L) in March but we missed the time period during April/May as the samples analyzed here were taken from October 2015 to March 2016. Another interesting observation at the Dhaka site was that the number of *ctxA* gene copies detected was always slightly higher than *rfb*O1 (∼20%). It is known from previous research that El Tor strain N16961 carries a single copy of the cholera toxin prophage whereas there could be variation in copy number in other El Tor *V. cholerae* strains arising from selective pressure (Chun et al., 2009; Davis et al., 1999; Mekalanos et al., 1983; Trucksis et al., 1998). Also, the presence of CTX phages in the aquatic environment may significantly impact the abundance of *ctxA* positive cells in the environment.

In contrast to the inland Dhaka site, the O1 serogroup was only detected at low abundance or was undetectable at the two coastal sites sampled in Bangladesh. *V. cholerae* O1 was found at very low abundance in Kuakata (May 2014) and absent from Mathbaria during the single month sampled (FIG 5 and FIG 6), whereas in a previous study, detection of toxigenic *V. cholerae* O1 by DFA (direct fluorescent antibody) in water samples collected bi-weekly from March to December 2004 from six ponds in Mathbaria fluctuated from < 10 to 3.4×10^7^ CFU/L (Alam et al., 2006). It is noteworthy that during the period of this latter study, cholera cases recorded in Mathbaria were due to *V. cholerae* O1. Moreover, in the same study, no strains belonging to the O1 serogroup were found among the six hundred strains isolated from coastal areas in Bangladesh, confirming our observations in an area (Oyster pond, MA, USA) non-endemic for cholera that *V. cholerae* O1 is challenging to isolate (TABLE 3). The constant presence of *V. cholerae* O1 in Dhaka as opposed to a more stochastic appearance in coastal locations suggests that the level of human interaction with water bodies influences its population dynamic.

### Higher abundance of *V. cholerae* and *V. metoecus* in larger size fractions indicates preference for particle association

Amongst the samples collected from endemic (Bangladesh) and non-endemic (Oyster Pond, USA) regions, overall counts of *V. cholerae* ranged from 1×10^4^ copies/L to 1×10^6^ copies/L for each sample (FIG 3, FIG 5 and FIG 7). In Oyster pond, most of *V. cholerae* (∼93%), including ∼18% of the O1 serogroup *V. cholerae*, were detected in the largest fraction size (>63 μm). Similarly, the majority of *V. cholerae* cells from coastal areas of Bangladesh (∼62%) were detected in the largest fraction size (> 63 μm) (FIG 5 and FIG 6). The toxigenic *V. cholerae ctxA* marker gene and O1 serogroup strains *rfbO1* gene were found only in the >63 μm size fraction and in very low concentrations in Kuakata (3.4×10^3^ and 2.9×10^3^ copies/L, respectively) (FIG 6).

This skewed distribution of *V. cholerae* toward the largest fraction size (FIG 4 and FIG 6) suggests that most cells are associated with large particles, zooplankton, or phytoplankton hosts in environmental reservoirs (Ali et al., 2012; Chun et al., 2009; Faruque et al., 1998; Meibom et al., 2004). This explains why in rural areas of Bangladesh, filtering environmental water through folded cloth facilitates the reduction of particles and debris and significantly lowers the risk of cholera (Rosenberg, 2009).

## CONCLUSION

This research describes for the first time a sensitive multitarget real-time qPCR application for the simultaneous detection and quantification of *V. cholerae* and *V. metoecus* from environmental water samples. The *viuB* and *mcp* markers are specific to *V. cholerae* and *V. metoecus*, respectively, and made it possible to quantify the absolute abundance of members of these two species in DNA extracted from environmental biomass. The *ctxA* and *rfbO1* markers are specific for detection of toxigenic and O1 serogroup strains, allowing determination of the proportion of the total *V. cholerae* population represented by these strains. This is also a fundamental study on quantification of *V. cholerae* on a significant scale in a cholera-endemic area. Although cholera has two seasonal peaks in this region, we showed that toxigenic *V. cholerae* O1 was persistent in the inland water reservoir at levels that pose a risk to human health. *V. metoecus* was not detected in this area, indicating a different geographical distribution than that of its closest relative. The sporadic presence of *V. cholerae* O1 at a substantial proportion of the local *V. cholerae* total population in a region not endemic for cholera also highlights the wide distribution of this lineage displaying potential for the emergence of novel virulent variants.

## Acknowledgement

MA of ICDDR, B thanks the government of Bangladesh, Canada, Sweden and United Kingdom for providing unrestricted core support. We would like to thank Norman Neumann, professor and vice-dean, school of public health, University of Alberta for reviewing the manuscript.

