## Supplementary materials for "Simultaneous quantification of *Vibrio metoecus* and *Vibrio cholerae* with its O1 serogroup and toxigenic subpopulations in environmental reservoirs"

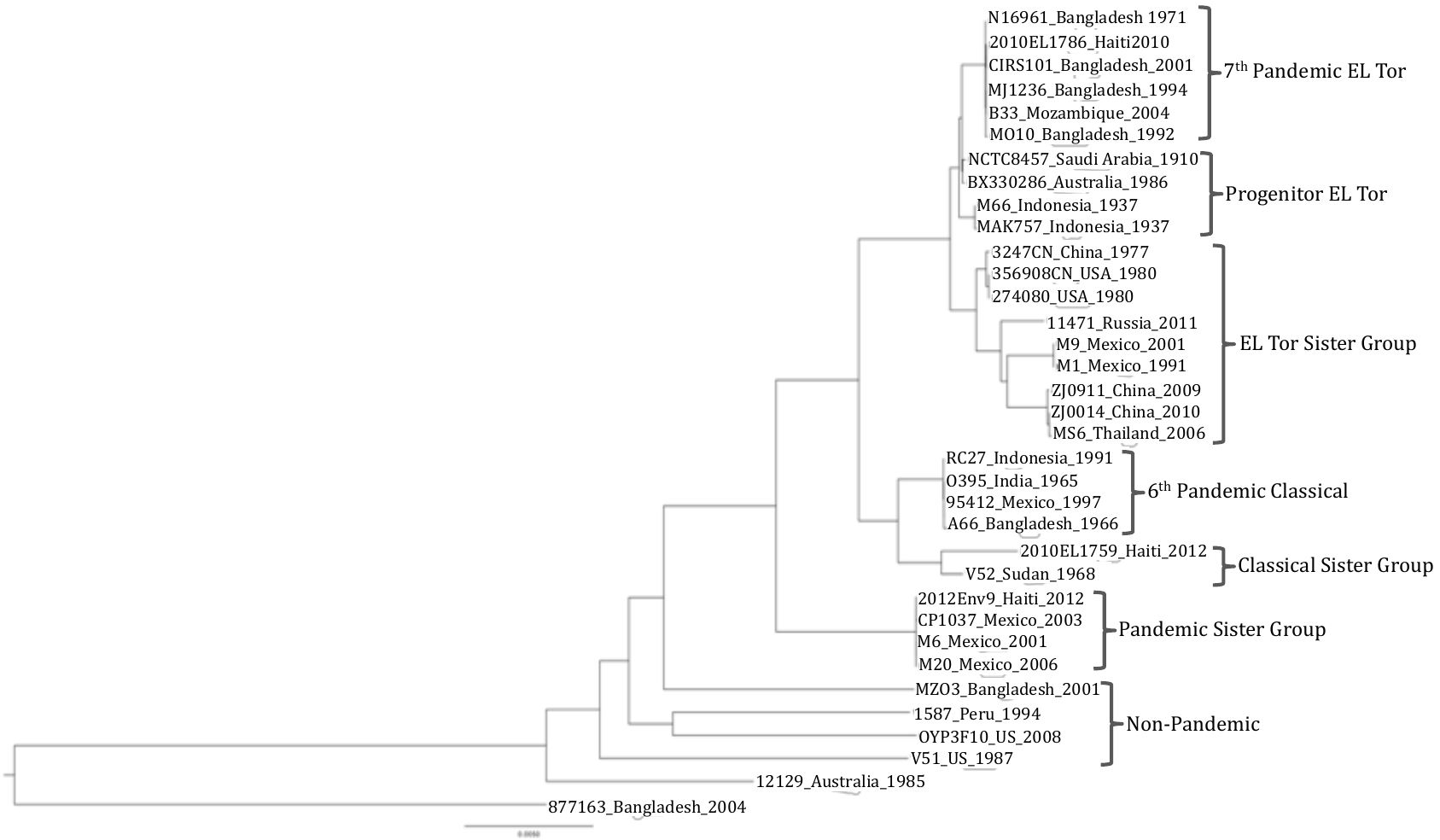

**FIG S1.** **Phylogeny of the pandemic generating lineage of *V. cholerae*.** The maximum likelihood phylogenomic tree was constructed from the alignment of locally collinear blocks (2,367,372 bp) using GTR gamma substitution model with 100 bootstrap replicates (all nodes had >98% bootstrap support). Environmental strain *Vibrio sp.* strain 877-163 was used to root the tree. The source and isolation year of the strains used are indicated next to the strain names.

**TABLE S1. Bacterial strains used for validating primers and probes in this study**

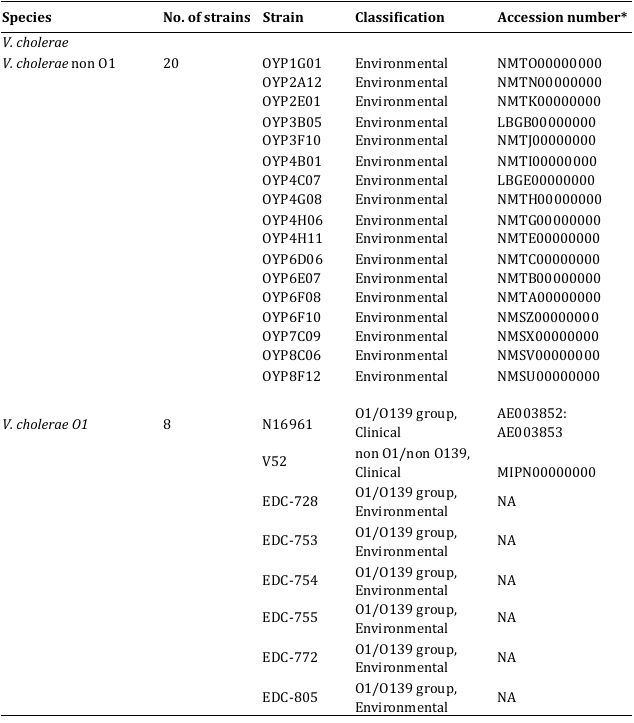

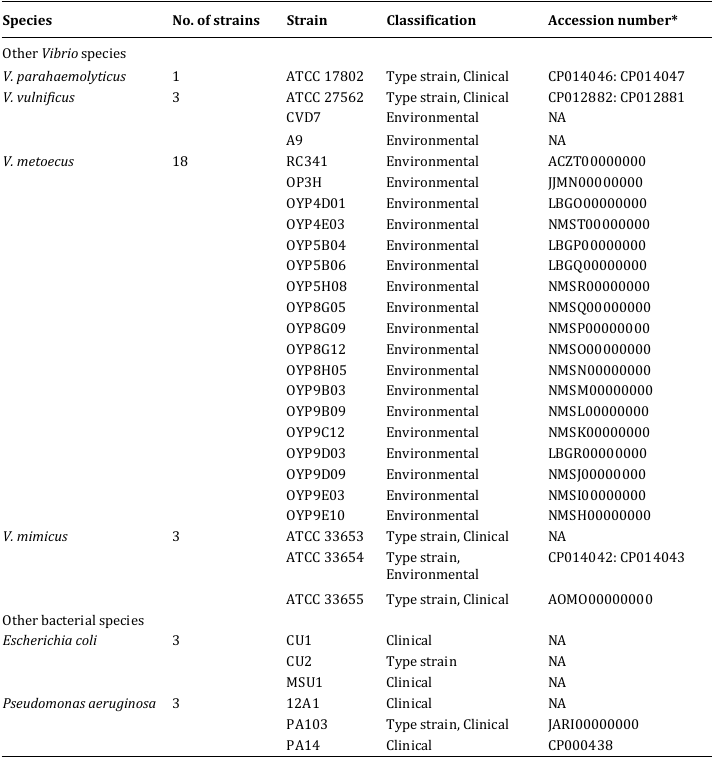

* Accession number obtained from Genebank

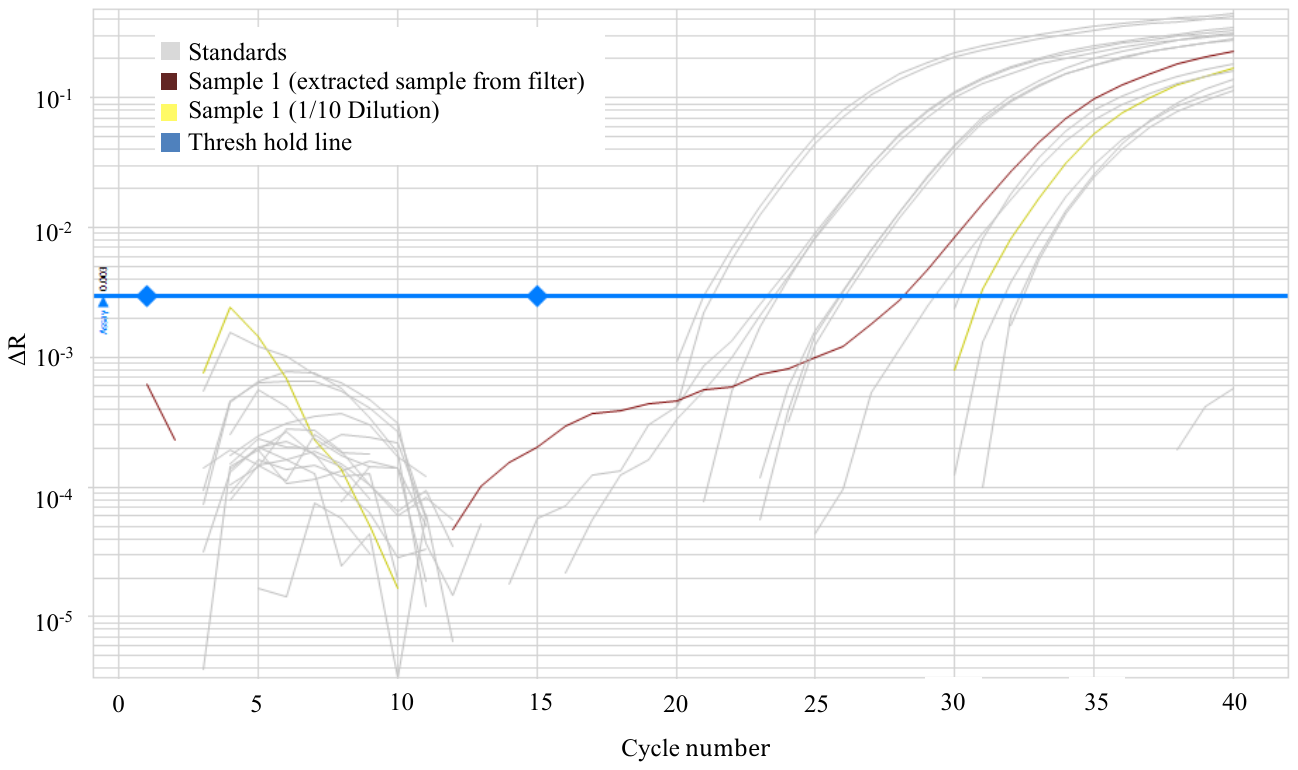
**FIG S2.** **Testing of inhibition in qPCR amplification.** The qPCR assay was performed on an extracted DNA sample treated with One step PCR inhibitory removal kit (sample1) from Oyster Pond large fraction size (> 63 μm) and a 10× dilution. Standards (3×10^4^ copies to 3 copies per reaction indicated with grey colored lines) were run in the same experiment. ΔR, normalized reporter value. The 10× dilution of samples shifted the Cq values by 3.3 cycles ± 0.05, indicating no inhibition.

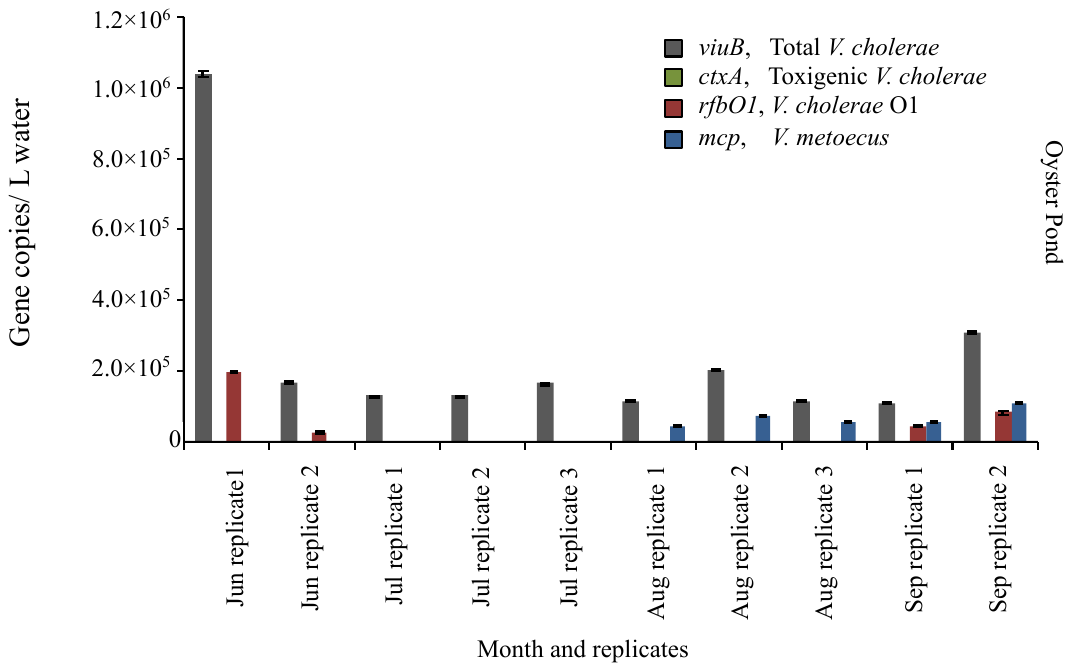

**FIG S3.** **Biological replicates of the quantification of *V. cholerae* along with its toxigenic and serogroup O1 subpopulations and its close relative *V. metoecus* in Oyster Pond, MA, USA*.*** Environmental water samples were collected in two or three replicates during the months of June to September in 2008. Each replicate represents 50 ml filtered independently from the same 10 L sample as its matching replicate. Bacteria were quantified by qPCR of marker genes. The *viuB* gene was used to quantify total *V. cholerae*; *ctxA* and *rfbO1* were used to measure toxigenic *V. cholerae* and *V. cholerae* O1, respectively; the abundance of *V. metoecus* was quantified using the *mcp* gene. Each qPCR reaction was run in triplicate. Mean values are shown with error bars indicating the standard deviation between technical replicates.
